# A VLP vaccine platform comprising the core protein of hepatitis B virus with N-terminal antigen capture

**DOI:** 10.1101/2024.06.07.597873

**Authors:** Kaniz Fatema, Joseph S. Snowden, Alexander Watson, Lee Sherry, Neil A. Ranson, Nicola J. Stonehouse, David J. Rowlands

## Abstract

Nanoparticle presentation systems offer the potential to develop new vaccines rapidly in response to emerging diseases, a public health need that has become increasingly evident in the wake of the COVID-19 pandemic. Previously, we reported a nanoparticle scaffold system termed VelcroVax. This was constructed by insertion of a high affinity SUMO binding protein (Affimer), able to recognise a SUMO peptide tag, into the major immunodominant region of VLPs assembled from a tandem (fused dimer) form of hepatitis B virus (HBV) core protein (HBc). Here we describe a modified form of VelcroVax, comprising monomeric HBc with the Affimer inserted at the N-terminus (termed N-VelcroVax). In contrast to the tandem form of VelcroVax, N-VelcroVax VLPs were expressed well in *E. coli.* The VLPs effectively bound SUMO-tagged Junín virus glycoprotein, gp1 as assessed by structural and serological analyses. Cryo-EM characterisation of N-VelcroVax complexed with a SUMO-Junín gp1 showed continuous density attributable to the fused Affimer, in addition to evidence of target antigen capture. Collectively, these data suggest that N-VelcroVax has potential as a versatile next generation vaccine scaffold.

## Introduction

Virus vaccines developed by conventional means, e.g. attenuation or chemical inactivation, have been highly effective in reducing disease burdens in both humans and domesticated animals. Furthermore, advances in molecular biology and immunology in recent decades have expanded the options available for development of novel vaccines against intractable diseases and against newly emerging pathogens. Newer approaches to vaccine development include recombinant expression of protein subunits, recombinant live viral vectors, DNA, mRNA and nanoparticle vaccine platforms [1–3]. The original nanoparticle vaccines were virus-like particles (VLPs) produced by recombinant expression of self-assembling viral structural proteins. The first VLP vaccine to be licenced for human use was the hepatitis B surface antigen vaccine (HBsAg), which has been used to immunise many millions of people since its introduction in the 1980s [4, 5]. More recently, VLP vaccines have been successfully developed to immunise against human papillomavirus. These VLP vaccines provide immunity against the viruses from which the VLPs originated [6]. However, the principle of using VLPs as presentation systems for antigens derived from heterologous pathogens has been exemplified by the licencing of Mosquirix, which comprises HBsAg modified to include antigenic components of the malaria parasite [7]. In addition to viral structural proteins, other self-assembling proteins have been shown to produce nanoparticles suitable for chemical modification to provide candidate vaccine platforms [8, 9].

VLPs lack viral genetic material and therefore have no replicative ability and are intrinsically safe. Additionally, due to their structural resemblance to native viruses, VLPs inherently possess “self-adjuvanting” immunogenic properties to elicit a strong immune response. Moreover, VLPs are able to elicit both humoral and cellular immune responses initiating the production of specific antibodies and activating cytotoxic T-cells [10–13]. In addition to acting as antigens themselves, VLPs can be modified to display foreign epitopes [14–16]. A key advantage of using VLP-based vaccine platforms is the ability to present antigens in a dense, highly repetitive manner, which leads to cross-linking of B cell receptors (BCRs) and B cell activation. Many studies have shown that VLP-based vaccines containing multivalent antigens elicit robust B cell activation and signalling and induce more effective immune responses than monovalent antigens [11, 13, 17, 18].

The capsid (or core) protein of hepatitis B virus (HBc) readily forms icosahedral VLPs when expressed in a wide range of prokaryotic and eukaryotic systems, which are inherently highly immunogenic. Consequently, the HBc VLP system has long been investigated as a potential vaccine platform [14, 15, 19–21]. HBc protein monomers comprise either 183 or 185 amino acids and dimerise to produce structural subunits which assemble to form icosahedral VLPs with *T =* 3 (90 dimers) or *T =* 4 (120 dimers) morphologies [22]. The dimeric assembly results in the formation of 4 helix bundles (spikes) which project from the particle surface and a sequence at the tips of the spikes comprises the major immunodominant region (MIR). The MIR has been used as a preferred site for insertion of foreign epitopes because of its immunodominance [16, 23, 24] but this also poses challenges. There is a potential for steric clashes when two closely located copies of the inserted sequence are present at the tips of the spikes [23, 25]. Furthermore, insertion of large or hydrophobic sequences can interfere with assembly and therefore abrogate VLP formation [26, 27]. We previously addressed this problem by fusing two copies of HBc protein to produce tandem HBc VLPs, which allows the insertion of a foreign sequence into one MIR while leaving the other MIR in the tandem partner unmodified, resulting in the presence of only one foreign epitope per spike [23]. This approach allowed us to develop VelcroVax, incorporating an antigen capture sequence, an Affimer (a non-antibody high affinity artificial binding protein), inserted at a single MIR per spike [16]. Although these constructs form VLPs when expressed in eukaryotic systems, such as yeast or plants, they do not assemble efficiently when expressed in *E.coli*. Consequently, here we have developed an alternative antigen capture platform compatible with expression in *E.coli*. The MIR is the most exposed part of the VLP and consequently is the immunodominant region [19, 23]. Therefore, in order to reduce the inherent anti-hepatitis B virus immunogenicity of the VLPs, we modified the natural HBc protein sequence by inserting 5 additional amino acids within the MIR. This construct (which we refer to as wild type (wt) HBc 190) was modified to include an Affimer fused to the N-terminus and termed N-VelcroVax. The resulting VLPs were able to bind a viral glycoprotein (Junín gp1) bearing the appropriate affinity tag as demonstrated by ELISA, gradient centrifugation and cryo-EM.

## Results

### Expression of N-VelcroVax in *E. coli*

We previously generated a vaccine platform system termed VelcroVax in which an anti-SUMO Affimer was inserted into the MIR of one of the HBc monomers within a fused HBc dimer. The Affimer was presented on the surface of assembled VLPs and was able to bind SUMO-tagged target antigens [16]. Here, we have modified a monomeric HBc protein containing a 5 amino acid insertion into the MIR, termed wt HBc 190 (Fig.1A), by introducing the anti-SUMO Affimer sequence (Fig. 1C) at the N-terminus to produce N-VelcroVax (Fig 1B). This protein was expressed in *E.coli*, using the ClearColi BL21 (DE3) strain to minimise contamination with bacteria-derived pyrogens. Both wt HBc 190 and N-VelcroVax were efficiently expressed, as demonstrated by Western Blot analysis using anti-HBc (10E11) antibodies (Fig. 1D), with proteins of 22 kDa and 35 kDa detected, as expected. A minor band apparent in the N-VelcroVax sample is likely to be a degradation product.

**Figure 1:**
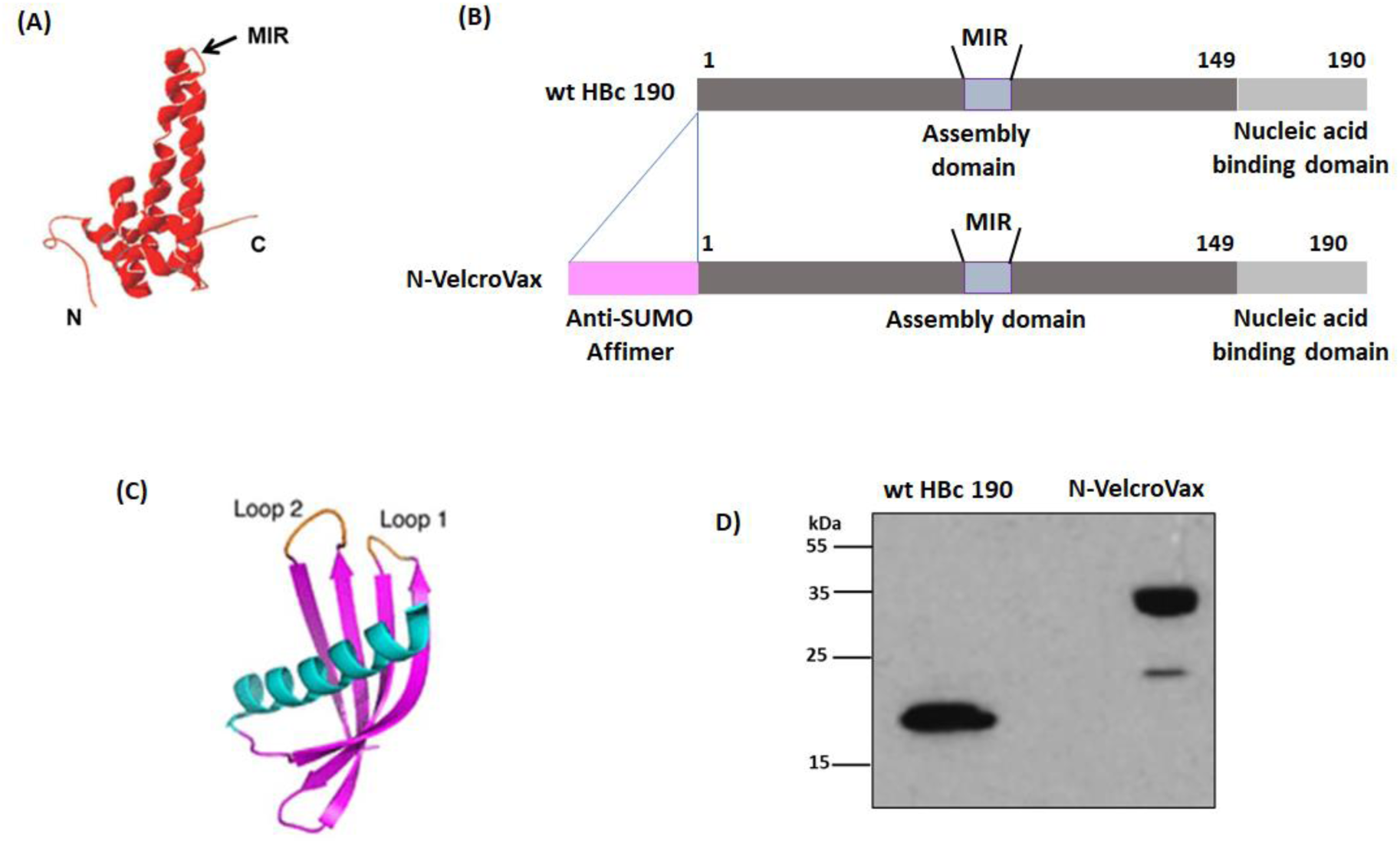
Characterisation of N-VelcroVax. **(A)** X-ray crystal structure of HBc monomer indicating the N-terminus, C-terminus and major immunodominant region (MIR) (PDB: 1QGT; ref. [48]). **(B)** wt HBc 190 construct (contains no anti-SUMO-Affimer) and N-VelcroVax, indicating the N-terminal insertion of the anti-SUMO-Affimer. **(C)** X-ray crystal structure of an Affimer selected against human SUMO protein (PDB: 5ELJ). Loop 1 and Loop 2 are the variable regions [49]. **(D)** Small-scale expression of HBc VLPs: Western Blot of wt HBc 190 and N-VelcroVax expressed in ClearColi BL21 (DE3) *E. coli* cells detected with mouse monoclonal anti-HBc (10E11). The figure is a representative example of three separate experiments.

### N-VelcroVax assembles into VLPs

After confirmation of expression of N-VelcroVax in ClearColi BL21 (DE3) *E. coli*, cell extracts were purified by differential ultra-centrifugation including separation of assembled VLPs on sucrose gradients. Gradient fractions were then analysed by Coomassie blue staining and Western blot which confirmed the presence of N-VelcroVax VLPs (Fig. 2A) [16]. The purity of the sucrose gradient separated VLPs can be assessed from the lack of visible contaminating proteins in the Coomassie stained gel. Examination of the peak fraction (fraction 5) by negative stain transmission electron microscopy (TEM) showed VLPs of the expected morphology (Fig. 2B).

**Figure 2.**
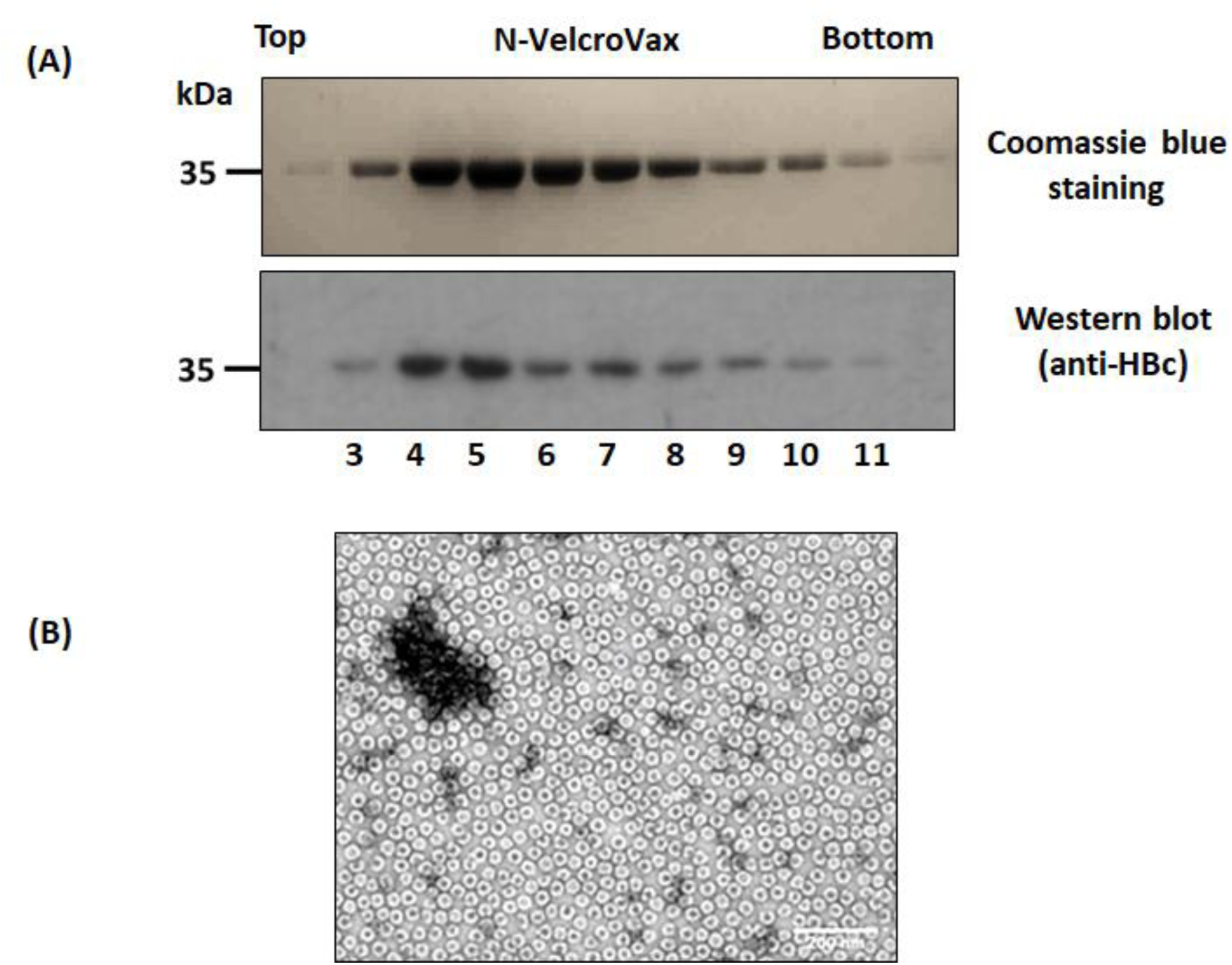
Purification and characterisation of N-VelcroVax particles. **(A)** Coomassie blue staining (upper panel) and Western blot (lower panel) of gradient purified N-VelcroVax particles, expressed in ClearColi BL21 (DE3) *E. coli* cells detected with mouse monoclonal anti-HBc (10E11). **(B)** Negative stain TEM analysis of N-VelcroVax. The VLPs were stained using 2% uranyl acetate. The scale bar shows 200 nm. The figure is a representative example of three separate experiments.

### Complexes of SUMO-Junín gp1 and N-VelcroVax VLP co-sediment during sucrose gradient centrifugation

We examined the SUMO-binding properties of N-VelcroVax VLPs using a SUMO-tagged gp1 protein of Junín virus, which we had previously shown to bind to the Affimer when presented at the MIR of tandem VelcroVax [16]. For the initial investigation of binding, N-VelcroVax VLPs were mixed with SUMO-Junín gp1 at a 1:1 molar ratio overnight at 4°C and separated by 15 - 45% sucrose gradient ultracentrifugation. The resulting fractions were analysed by Western blot using anti-HBc (10E11) and anti-Junín antibodies to detect the positions of N-VelcroVax (Fig. 3A) and SUMO-Junín gp1 (Fig. 3B) respectively in the gradients. Both HBc N-VelcroVax (35 kDa) and SUMO-Junín gp1 (54 kDa) were present in the same gradient fractions, indicative of a non-covalent interaction (Fig. 3). As shown in Fig. 3A, the majority of the SUMO-Junín gp1 co-sedimented with the VLPs, although some unbound material (29.8 +/- 3.8 %) was detected in the top gradient fractions.

**Figure 3.**
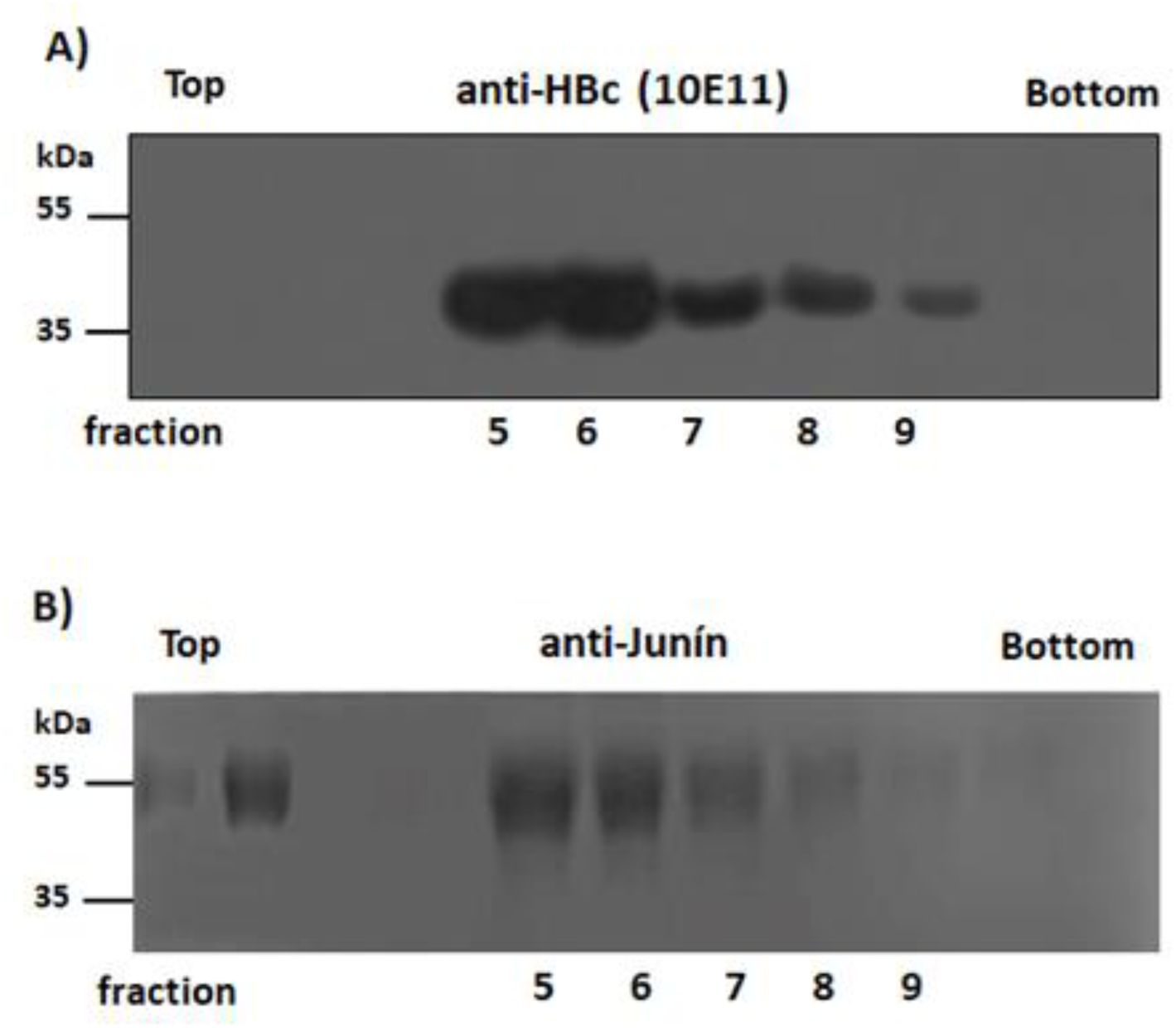
Evaluating the binding of N-VelcroVax to SUMO-Junín gp1. N-VelcroVax and SUMO-Junín gp1 were mixed and incubated overnight before separation on a sucrose density gradient. Gradient fractions were analysed by Western blot using either mouse monoclonal anti-HBc (10E11, A) or anti-Junín gp1 (NR 2567, B). The figure is a representative example of three separate experiments.

### Evaluation of SUMO-Junín gp1 binding to N-VelcroVax by ELISA

The SUMO-Junín gp1 interaction with HBc N-VelcroVax was assessed by ELISA. SUMO-Junín gp1 and N-VelcroVax were mixed before loading onto ELISA plates. Molar ratios ranged from 1:1 to 5:1 (HBc monomer:gp1). After washing, the wells were interrogated with anti-HBc or anti-SUMO-Junín gp1 antisera. The results indicated that N-VelcroVax VLPs clearly bound SUMO-Junín gp1 with maximum binding occurring when the components were mixed at a 1:1 ratio (Fig. 4). No binding of SUMO-Junín gp1 to wt HBc 190 VLPs was detected, as expected (Fig. 4).

**Figure 4.**
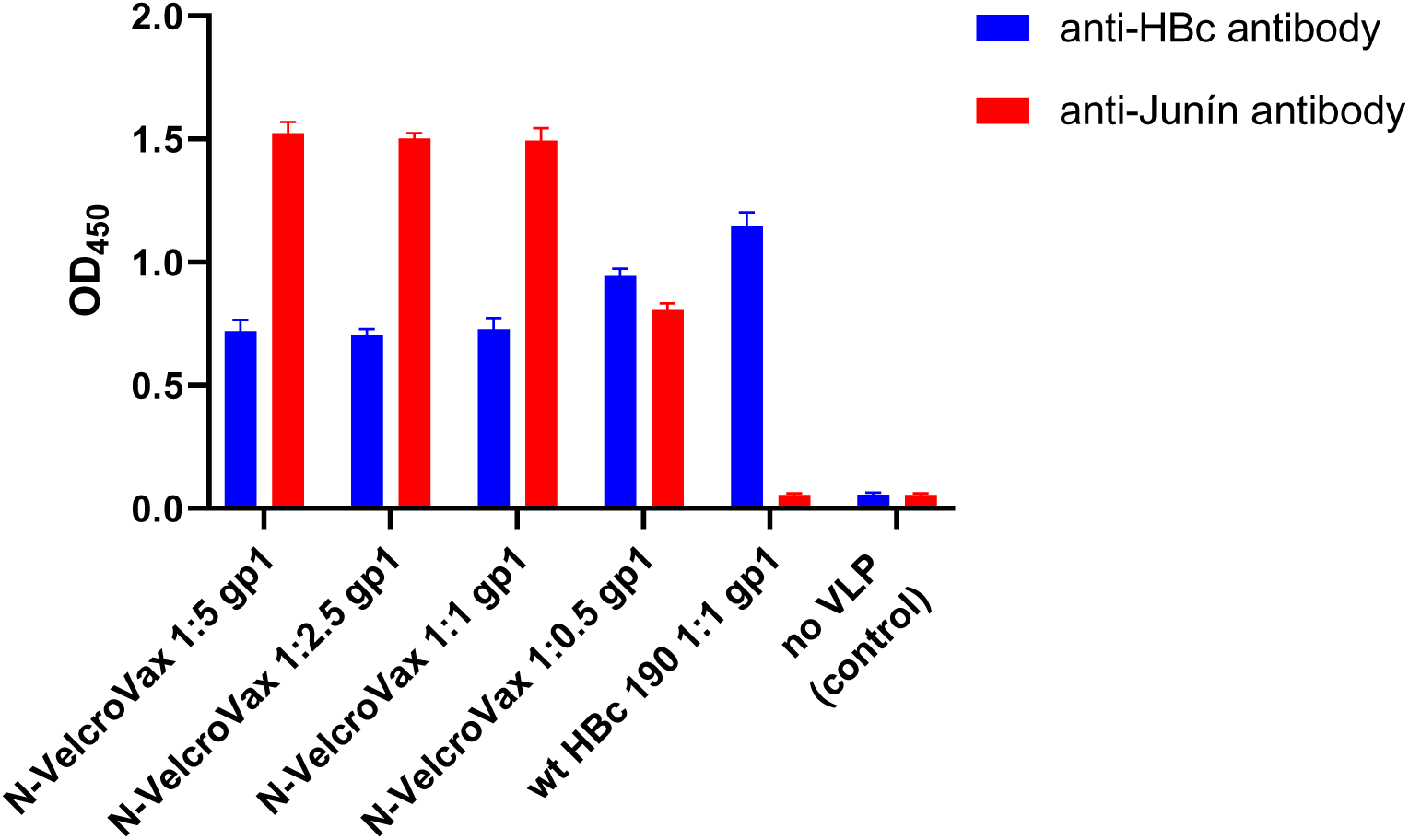
Evaluation of the interaction between N-VelcroVax and SUMO-Junín gp1 by ELISA. N-VelcroVax and SUMO-Junín gp1 were mixed in molecular ratios ranging from 1:1 to 1:5 and binding determined by ELISA. Anti-HBc 10E11 (1:1000) and anti-Junín gp1 NR 2567 (1:32,000) were used to detect HBc VLPs and SUMO-Junín gp1 respectively. Anti-mouse antibody conjugated with HRP was added as secondary antibody. TMB chromogenic substrate was used to detect HRP. The optical density at 450 nm (OD 450 nm) is represented in arbitrary units (*n* = 3).

### Structural characterisation of N-VelcroVax and N-VelcroVax:SUMO-Junín gp1 particles

N-VelcroVax samples were vitrified for cryo-EM data collection (Fig. S1, Table S1). This confirmed two different sizes of icosahedral particles. Reconstructions of each particle stack with icosahedral symmetry imposed revealed that these corresponded to *T =* 3 and *T =* 4 configurations, as expected.

Interestingly, both reconstructions (resolved to ∼3.5 Å and ∼3.3 Å, respectively) showed continuous densities located between the four-helix bundles on the surface of the VLPs (Fig. 5). Each of these densities was of appropriate dimension to accommodate an Affimer and was located close to the residue at the N-terminus of a wt HBc 190 monomer, as expected. Unstructured internal density was also observed, possibly representing non-specific binding of nucleic acid derived from the *E.coli* expression system as reported previously [28] (Fig. 5).

**Figure 5.**
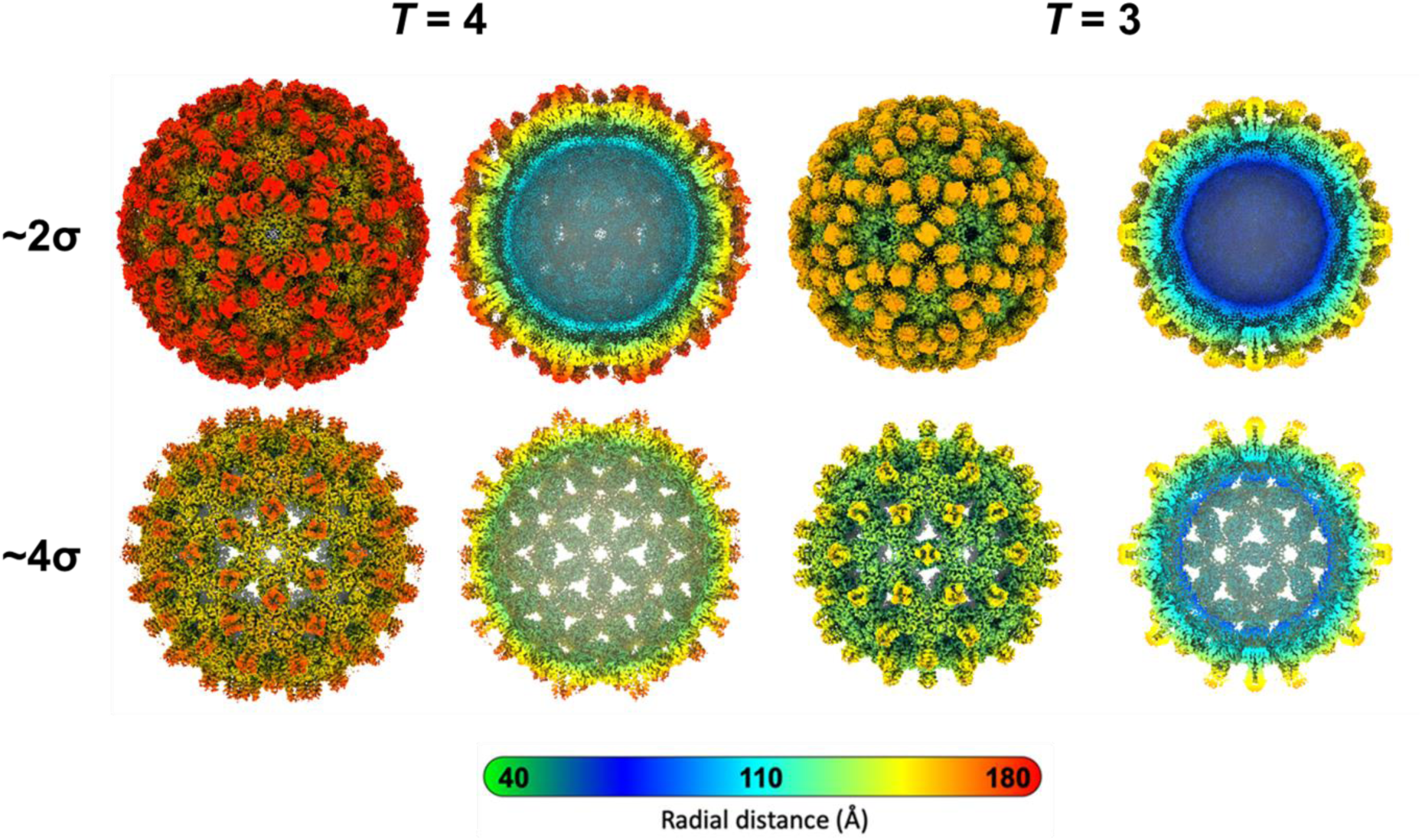
Characterisation of N-VelcroVax VLPs by cryo-EM. Density maps for *T =* 4 (left) and *T =* 3 (right) N-VelcroVax reconstructions, filtered according to local resolution. Each density map is shown as a complete isosurface representation and central cross-section at both low (approx. 2σ, Affimer densities visible) and high (approx. 4σ, Affimer densities not visible) contour level. Cross-sectional views show density, potentially nucleic acid bound non-specifically to the interior surface of the VLPs. Maps are coloured by radial distance (Å).

Next, we attempted to visualise N-VelcroVax VLPs in complex with a SUMO-tagged antigen, specifically, the gp1 glycoprotein of Junín virus (SUMO-gp1). As initial screening by negative stain TEM of N-VelcroVax:SUMO-gp1 complexes revealed significant aggregation, an alternative on-grid binding approach was pursued (i.e. separate samples of N-VelcroVax and SUMO-gp1 were applied directly to the grid in sequence). This showed no evidence of aggregation of N-VelcroVax particles and visual inspection suggested that the boundaries of the VLPs had become less well defined, indicative of decoration with SUMO-gp1 molecules (Fig. 6A).

**Figure 6.**
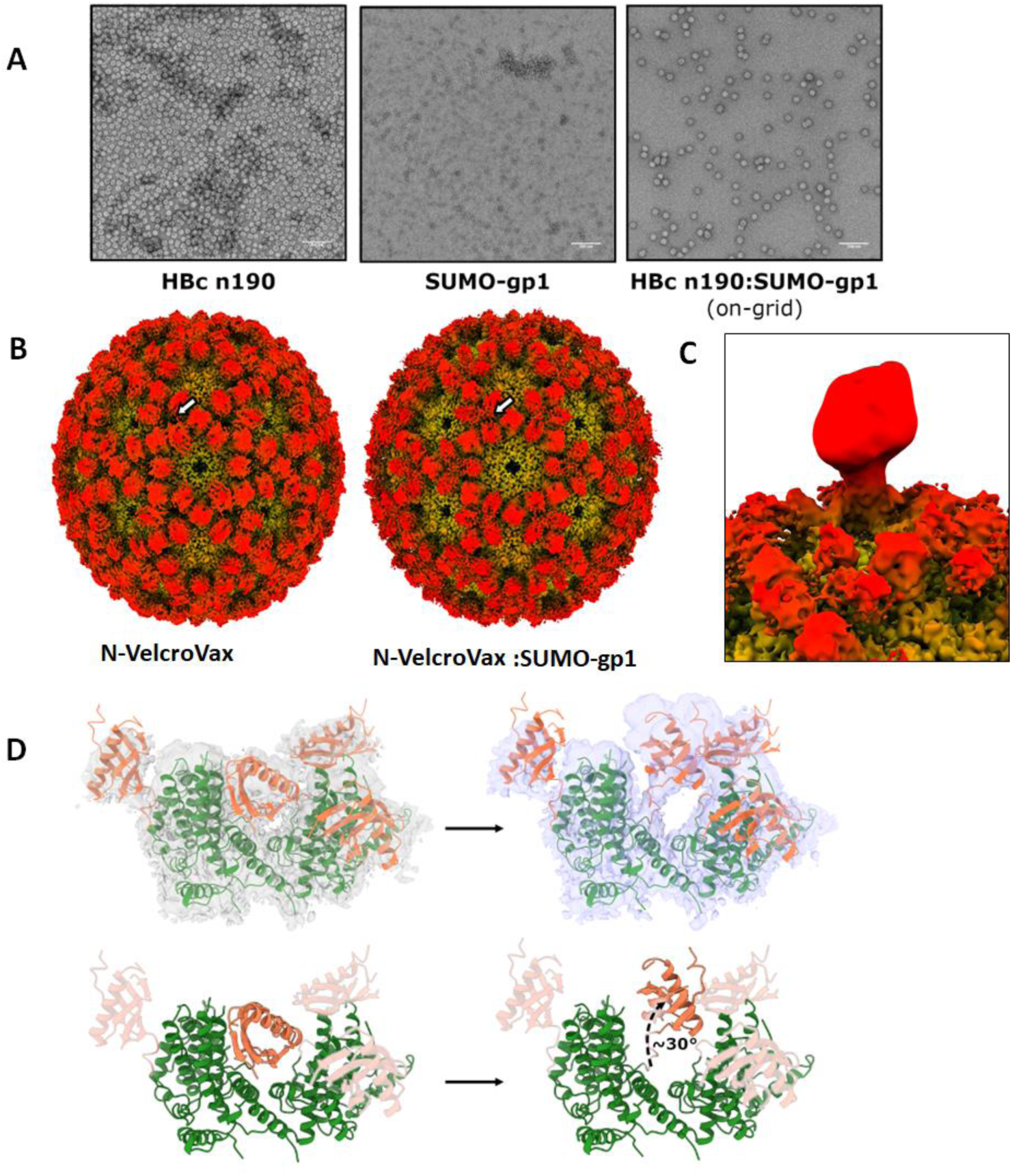
Structural characterisation of N-VelcroVax in complex with SUMO-gp1. **(A)** Representative micrographs from negative stain TEM analysis of N-VelcroVax alone (left), SUMO-gp1 alone (centre), or N-VelcroVax:SUMO-gp1 following on-grid interaction (right). Scale bars show 200 nm. **(B)** Isosurface representations of cryo-EM density maps for the *T =* 4 configuration of unliganded N-VelcroVax (left) and N-VelcroVax:SUMO-gp1 (right), shown at ∼2 σ and coloured radially. Affimers show a subtle change in relative positioning, as indicated by the white arrow. **(C)** Asymmetric reconstruction of particles from a single focussed class following focussed classification of N-VelcroVax:SUMO-gp1 *T =* 4 particles, showing additional low-resolution density corresponding to bound SUMO-gp1. All focussed classes are shown in Fig. S3. **(D)** Density from an asymmetric reconstruction of *T =* 4 N-VelcroVax:SUMO-gp1 following focussed classification (purple) is suggestive of a reorientation of the Affimer compared to its position in the unliganded N-VelcroVax density map (grey). Fitted atomic models (orange – Affimers, green – remainder of N-VelcroVax) are shown for illustrative purposes.

This on-grid binding approach was subsequently used to prepare samples for cryo-EM analysis of N-VelcroVax:SUMO-gp1 (Fig. S2, Table S1). Processing of cryo-EM data yielded density maps for *T =* 3 and *T =* 4 VLPs resolved to 3.0 Å and 3.5 Å, respectively. Initial inspection of density maps revealed no additional density attributable to bound SUMO-gp1, although there appeared to be a subtle change in the orientations of the Affimers, with these being closer together in the N-VelcroVax:SUMO-gp1 complex map compared to the unliganded N-VelcroVax map (Fig. 6B).

Hypothesising that the SUMO-gp1 was perhaps bound at a low occupancy, we attempted to resolve density for the glycoprotein by performing focussed classification, as described previously [29–32]. A mask was applied to a single Affimer including the volume expected to contain SUMO-gp1 for both *T =* 3 and *T =* 4 VLPs (Fig. S3). Interestingly, several focussed classes revealed low resolution density continuous with Affimer density, suggesting that SUMO-gp1 had been captured by N-VelcroVax Affimers but was only present at a low occupancy (Fig. 6C), in contrast to the ELISA and sucrose density gradient data, which suggested that the occupancy by SUMO-gp1 was higher. This is most likely due to the marked differences in the methods used for complex formation for the solution-based and cryo-EM studies. Interestingly, density corresponding to the SUMO-gp1-binding Affimer in asymmetric reconstructions of focussed classes had reoriented upwards, suggesting that movement of the Affimer was necessary to accommodate the size of SUMO-gp1 (Fig. 6D).

## Discussion

The development of vaccine platforms suitable for multimeric presentation of antigenic components of important pathogens, so maximising their immune recognition, is an attractive approach toward novel vaccine development. To this end, we produced recombinant HBc VLPs (N-VelcroVax) with an anti-SUMO Affimer genetically fused at the N-terminus of HBc190 protein. Although HBc VLPs have been expressed in a number of expression systems, e.g., insect cells, mammalian cells, plants, yeast [6, 11, 13, 24, 33], using low cost expression systems would make the VLPs accessible more widely, especially for lower-to-middle income countries (LMICs). Hence, we opted for a cost-effective *E. coli* expression system which offers simple, fast and inexpensive production, ease of in-process control and high productivity [6, 13, 33–36]. However, contamination with high levels of endotoxin derived from the *E. coli* cell wall is a major drawback of this expression system [35, 37–39]. To address this problem, we used ClearColi BL21 (DE3) *E. coli* cells for expression of N-VelcroVax. A previous study showed that using ClearColi BL21 (DE3) *E. coli* for expression of apolipoprotein A and heat shock protein 70 significantly reduced endotoxin level as measured by LAL assay (10 μg/mL) and did not trigger endotoxic responses in HEK-Blue hTLR4 cells [37]. The endotoxin content of N-VelcroVax expressed in ClearColi BL21 (DE3) *E. coli* was 10.2 EU/mL as assessed by LAL assay, which is half the maximum recommended level of endotoxin acceptable for subunit vaccines [35, 37, 40].

Examination by negative stain TEM showed that the morphology of the N-VelcroVax VLPs was as expected [41, 42]. Cryo-EM analysis of the VLPs revealed particles in both *T =* 3 and *T =* 4 configurations, as expected. In our earlier analyses, Affimer density was not visible, likely as a result of the location of the Affimers on flexible linkers at the tips of four-helix bundles at the MIR [16]. However, for N-VelcroVax, in which Affimers are attached at the N-terminus and spatially constrained between the four-helix bundles, Affimer density was resolved. This suggests that the Affimers in the N-VelcroVax system are more rigidly constrained in their position compared to when presented at the MIR region.

N-VelcroVax was also visualised in complex with SUMO-gp1. Although SUMO-gp1 density was not resolved in symmetric reconstructions, focussed classification revealed low resolution density continuous with Affimer density. This finding is suggestive of low occupancy, with the majority of unoccupied sites leading to the ‘averaging out’ of SUMO-gp1 density in symmetric reconstructions. Low occupancy is likely a result of the on-grid binding method used during grid preparation to prevent particle aggregation. Indeed, both sucrose density gradient and ELISA assays demonstrated a much more efficient interaction in solution. Interestingly, an asymmetric reconstruction derived from particles belonging to a single focussed class revealed that the corresponding Affimer had undergone a major reorientation in order to accommodate the target antigen, indicative of some steric hindrance.

## Conclusion

Together with our earlier report of Affimer presentation at the MIR, these data demonstrate the flexibility of the HBc VLP system as a vaccine scaffold and open the possibility of a dual presentation system using both the N-terminus and the MIR for antigen display.

## Materials and methods

### Expression constructs

An anti-SUMO-Affimer sequence was genetically fused at the N-terminus of wt HBc 190 to produce N-VelcroVax (Fig. 2). This was subcloned into the pET29b expression vector using *Nde*1 and *Xho*1 and transformed into *E. coli* DH5α chemically competent cells for plasmid propagation.

For the target protein, Junín gp1, a SUMO-His6 tag [16] was genetically fused to the N-terminus of the Junín virus glycoprotein 1 to form SUMO-Junín GP1 (kindly supplied by Thomas A. Bowden, University of Oxford, UK). This protein was expressed in mammalian cells.

### Expression and purification of N-VelcroVax

ClearColi BL21 (DE3) *E. coli* was transformed with the N-VelcroVax plasmid and used for protein expression. N-VelcroVax plasmid was mixed with ClearColi BL21 (DE3) *E. coli* cells and incubated on ice for 30 min before being transferred to a 42°C heat block for 50 seconds. Heat-shock transformed cells were then plated on LB-Agar plates (30 μg/mL kanamycin) and incubated overnight at 37°C. A single colony was picked and cultured in 10 mL starter LB culture (30 μg/mL kanamycin) overnight at 37°C and shaking at 200 rpm. The preculture was diluted 1:1000 into 500 mL LB (30 μg/mL kanamycin) and cultured at 37°C, 200 rpm until OD_600_ ∼0.6 – 0.8. Protein expression was induced with isopropyl β-D-1-thiogalactopyranoside (IPTG) (0.1 μM) and incubated at 18°C, 200 rpm for a further 7 hours. The culture was centrifuged at 4000 × *g* for 10 minutes and the pellet was lysed with chemical lysis buffer. N-VelcroVax VLPs were purified by sedimentation through a 30% sucrose cushion followed by 15-45% sucrose gradient ultracentrifugation. Fractions (1 mL) were collected manually (top down) and fractions containing the VLPs were analysed by SDS-PAGE 12% gel where the protein bands were visualised by Coomassie blue R250 staining. The presence of HBcAg-reactive proteins was assessed by Western blot with MAb 10E11 using standard protocols. The protein content of fractions was assessed directly by BCA assay (Pierce, ThermoFisher Scientific).The concentration of purified N-VelcroVax was measured using bicinchoninic acid (BCA) assay (Pierce) which revealed a yield of protein ∼ 6.5 mg/L and endotoxin content of 10.2 EU/mL (Pierce LAL Chromogenic Endotoxin Quantitation kit; Thermo Scientific). Purified N-VelcroVax VLPs were stored at 4°C.

### Enzyme-linked immunosorbent assay (ELISA)

A non-competitive indirect ELISA was developed to assess the binding of N-VelcroVax vaccine scaffold to capture SUMO-tagged proteins using 96-well ELISA plates (Greiner Technologies, Bio-One, UK). The plate was coated with N-VelcroVax at 2.5 µg/mL (125 ng/well) and incubated overnight at 4 °C. The plate was blocked with blocking buffer (PBS + 2% skimmed milk) for 1 h at room temperature (RT) and then incubated with SUMO-tagged Junín gp1 for 1 hour at RT. Specific primary antibodies (mouse monoclonal anti-HBc 1 (10E11, Abcam ab8639), anti-Junín gp1 (NR 2567)) were added and incubated for 1 hour at RT. The bound primary antibodies were detected with HRP-conjugated goat anti-mouse IgG and incubated for 1 hour at RT. Subsequently, chromogenic substrate TMB (ThermoFisher, Waltham, MA, USA) was added and incubated for 20 minutes at RT. The plates were washed with wash buffer (PBS + 1% Tween 20) four times between each step. The reaction was stopped by 2 M H_2_SO_4_ (50 µL/well) and plates then analysed using a Biotech PowerWave XS2 plate reader.

### Negative-stain electron microscopy

Samples were prepared for negative-stain transmission EM by application to carbon-coated 300-mesh copper grids (Agar Scientific, UK) that had been glow discharged in air at 10 mA for 30 seconds immediately before use. Where N-VelcroVax or SUMO-Junín gp1 was imaged alone, 3 µL of sample was applied directly to the grid surface for 30 seconds before washing and staining. To probe the interaction of N-VelcroVax with SUMO-Junín gp1, the ligand was applied directly to the grid surface for 30 seconds after first applying N-VelcroVax and blotting away excess fluid. Following sample application, excess fluid was wicked away before grids were washed twice with 10 µL dH_2_O. To stain the sample, 10 µL 1-2% uranyl acetate solution was applied to the grid and immediately removed by blotting, before an additional 10 µL 1-2% uranyl acetate solution was applied for 30 seconds. Grids were blotted to leave a thin film of stain and left to air dry.

Imaging was performed using an FEI Tecnai F20 transmission EM (operating at 200 kV with a field emission gun), equipped with an FEI CETA camera (Astbury Biostructure Laboratory, University of Leeds). Data were collected at various defocus values (−2.0 µm to −5.0 µm) at a nominal magnification of 25,000× giving an object sampling of 0.418 nm/pixel.

### Cryo-electron microscopy

N-VelcroVax (in the presence or absence of target antigen) was vitrified using a LEICA EM GP plunge freezing device (Leica Microsystems, Wetzlar, Germany) in preparation for imaging by cryo-EM. Lacey carbon 400-mesh copper grids coated with a <3-nm continuous carbon film (Agar Scientific, UK) were glow discharged in air at 10 mA for 30 seconds prior to the application of 3 µL N-VelcroVax (or N-VelcroVax:SUMO-gp1). For N-VelcroVax:SUMO-gp1, an on-grid interaction approach was used, such that following the application of VLP, excess fluid was manually blotted away and 3 µL SUMO-gp1 was applied. The sample was incubated on the grid surface for 30 seconds at 80% relative humidity (8°C) before excess fluid was blotted for a duration between 1 – 4 seconds and the grid vitrified in liquid ethane (cooled to −179°C by liquid nitrogen). Grids were transferred to liquid nitrogen for storage prior to clipping and imaging with an FEI Titan Krios TEM (Astbury Biostructure Laboratory, University of Leeds). Data collection was performed in integrating mode at a magnification of 75,000× (calibrated object sampling of 1.065 Å/pixel) with the microscope operating at 300 kV (a full set of data collection parameters is provided in Table S1).

### Image Processing

Image processing was performed in Relion-3.0 and Relion-3.1 [43–45]. Following motion correction and CTF estimation of micrographs [45], a subset of manually picked particles was subjected to 2D classification, and class averages were used as templates for automated particle picking. Initial particle stacks generated by automated picking were down-sampled (2× for N-VelcroVax, 4× or 5× for N-VelcroVax:SUMO-gp1) before several rounds of 2D classification to progressively remove junk particles and separate *T =* 3 and *T =* 4 VLPs. To reduce computational load, the first 2D classification was performed with CTFs ignored until the first peak, and maximum signal limited to 200. *T =* 3 and *T =* 4 particles were separately re-extracted without down-sampling for independent 3D refinement (with I1 symmetry imposed) of each VLP configuration, based on initial models generated *de novo* in Relion. For each reconstruction, map quality was improved with iterations of CTF refinement and Bayesian polishing. Final refinements were performed with a solvent-excluding mask and flattened Fourier shell correlation (FSC) calculations. This was followed by mask-based sharpening, determination of nominal resolution using the gold-standard FSC criterion (FSC = 0.143), and calculation of local resolution for each map.

Focussed classification was performed using 2× down-sampled data to limit the computational load, following a previously described protocol [29]. Briefly, a cylindrical mask was generated in SPIDER [46] and placed manually over a single Affimer and the space distal to it, expected to contain density for SUMO-gp1, using UCSF Chimera [47]. Relion was used to apply a soft edge to the mask, and to generate symmetry-expanded stacks of particles based on the orientational information generated during separate symmetrised 3D refinements for *T =* 3 and *T =* 4 VLPs. Symmetry expanded particle stacks were subjected to 3D classification without alignments, with the focussed mask applied. Asymmetric reconstructions were performed using Relion following focussed classification.

## Acknowledgments

We thank other members of the Stonehouse/Rowlands group at the University of Leeds for their insightful contributions. We also thank Professor Tom Bowden and Dr Guido Paesen, University of Oxford who provided SUMO-Junín gp1.

## Authors and Contributions

K.F., J.S.S., N.J.S., and D.J.R. conceived and designed the experiments. K.F. and L.S. generated the N-VelcroVax sequence. K.F. introduced this into ClearColi BL21 (DE3) cells, generated material for the characterisation of N-VelcroVax and performed serological characterisation. J.S.S. performed negative stain EM and cryo-EM. J.S.S. and A.W. processed and analysed cryo-EM data. N.A.R., D.J.R. and N.J.S. provided supervision. K.F., J.S.S., L.S., D.J.R., & N.J.S. wrote and edited the manuscript. Funding was secured for this research by N.A.R and N.J.S.

---

**Figure S1.**
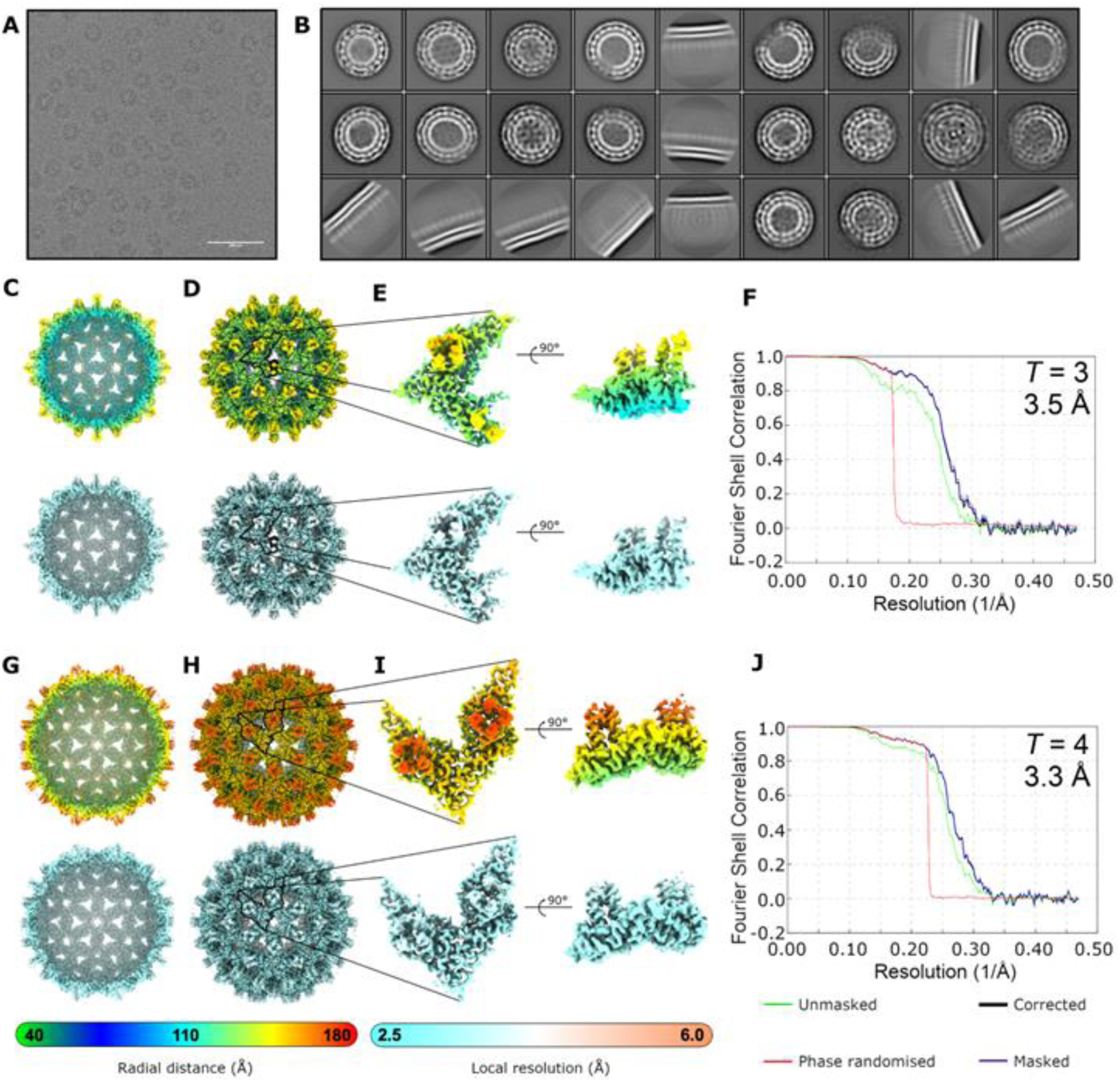
Processing of N-VelcroVax cryo-EM dataset. (A) Representative micrograph from N-VelcroVax data collection. Scale bar shows 100 nm. (B) 2D classes containing the most particles following automated particle picking. Data were two-fold down-sampled prior to classification. (C-E) Density map for *T =* 3 VLP of N-VelcroVax filtered according to local resolution, including (C) central section (D) whole VLP and (E) enlarged views of the asymmetric unit. (F) FSC plot for *T =* 3 N-VelcroVax reconstruction. (G-I) Density map for *T =* 4 VLP of N-VelcroVax. (J) FSC plot for *T =* 4 N-VelcroVax reconstruction. All reconstructions shown at ∼4 σ and coloured according to radial distance or local resolution, as indicated.

**Figure S2.**
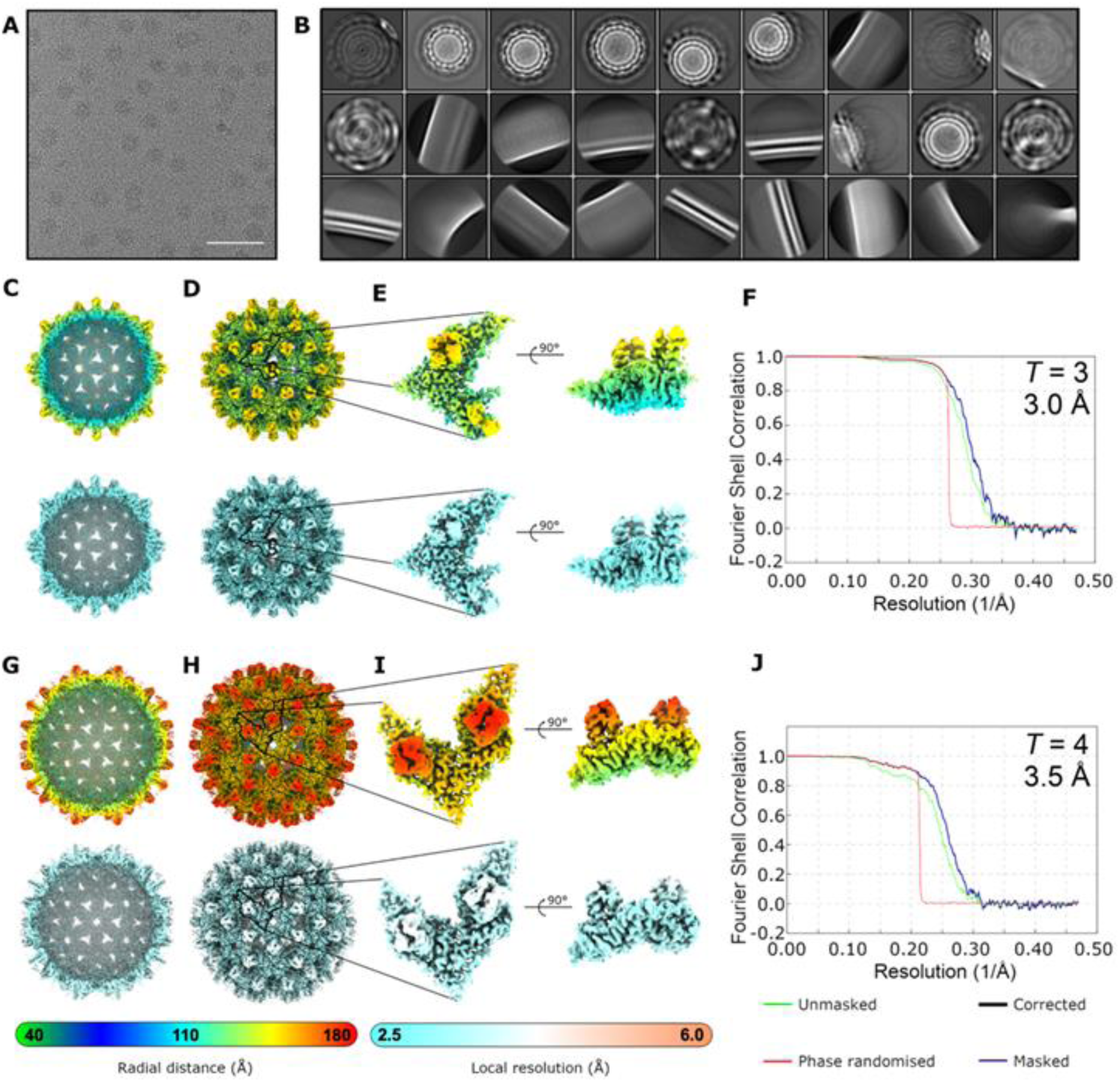
Processing of N-VelcroVax:SUMO-gp1 cryo-EM dataset. **(A)** Representative micrograph from N-VelcroVax:SUMO-gp1 data collection. Scale bar shows 100 nm. **(B)** 2D classes containing the most particles following automated particle picking. Data was five-fold down-sampled prior to classification. **(C-E)** Density map for *T =* 3 VLP filtered according to local resolution, including **(C)** central section **(D)** whole VLP and **(E)** enlarged views of the asymmetric unit. **(F)** FSC plot for *T =* 3 N-VelcroVax:SUMO-gp1 reconstruction. **(G-I)** Density map for *T =* 4 VLP. **(J)** FSC plot for *T =* 4 N-VelcroVax:SUMO-gp1 reconstruction. All reconstructions shown at ∼3 σ and coloured according to radial distance or local resolution, as indicated.

**Figure S3.**
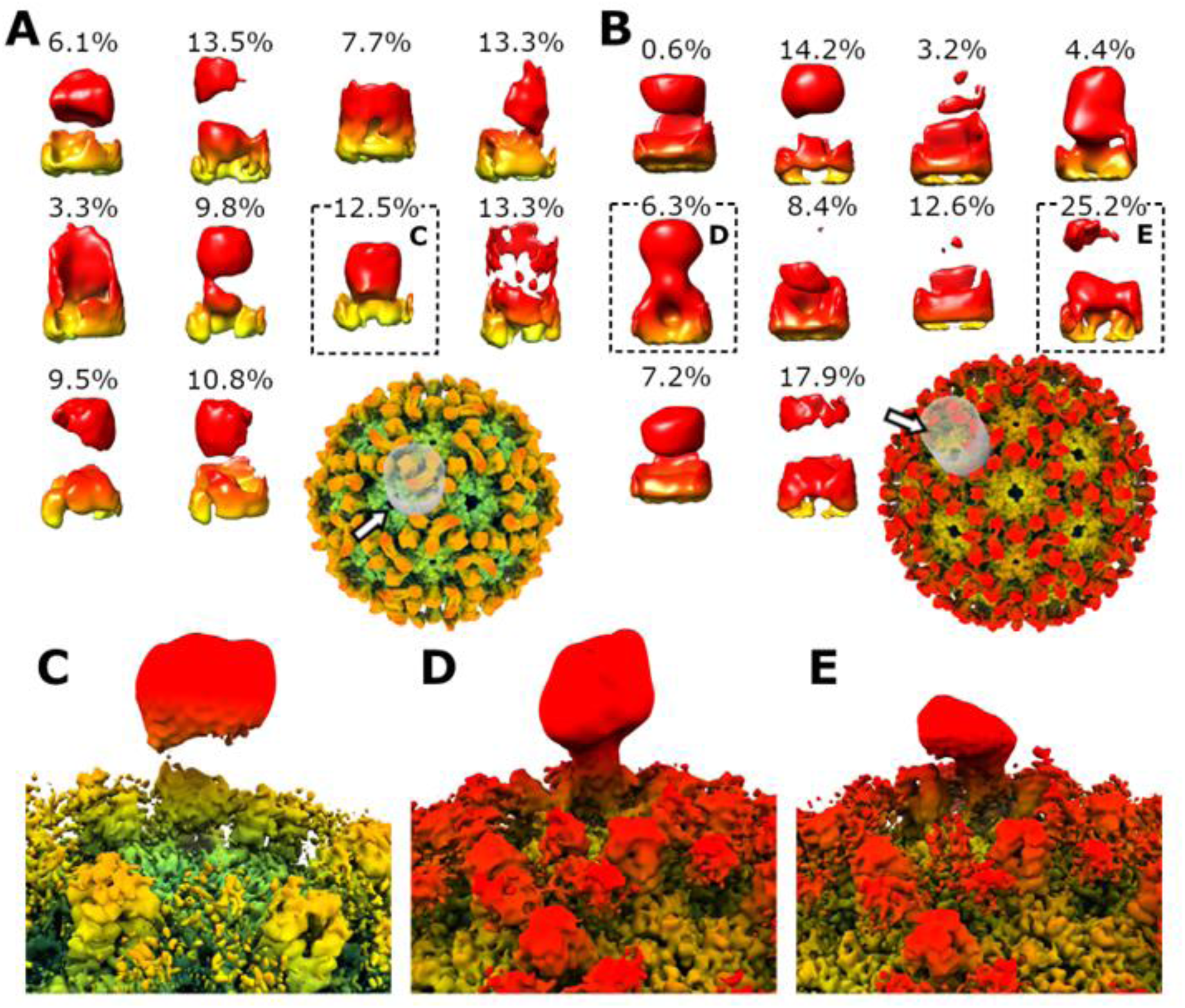
Focussed classification of N-VelcroVax:SUMO-gp1 cryo-EM dataset. **(A, B)** Density observed in all 10 focussed classes from focussed classification of **(A)** *T =* 3 N-VelcroVax:SUMO-gp1 and **(B)** *T =* 4 N-VelcroVax:SUMO-gp1. The number above each class indicates the proportion of sub-particles that were assigned. The position of the mask (grey) is shown for reference. Classes are shown oriented from the viewpoint indicated by the white arrows. **(C-E)** Asymmetric reconstructions using particles contained in the focussed classes indicated by dashed boxes in (A) and (B), filtered by local resolution.

**Table S1:** N-VelcroVax and N-VelcroVax:SUMO-Junín gp1 cryo-EM data collection and processing parameters.

| Sample | N-VelcroVax |  | N-VelcroVax:SUMO-gp1 |  |
| --- | --- | --- | --- | --- |
| Microscope | FEI Titan Krios |  | FEI Titan Krios |  |
| Detector mode | Linear |  | Linear |  |
| Camera | Falcon III |  | Falcon IV |  |
| Voltage (kV) | 300 |  | 300 |  |
| Pixel size (Å) | 1.065 |  | 1.065 |  |
| Nominal magnification | 75,000× |  | 75,000× |  |
| Exposure time (s) | 1.0 |  | 1.3 |  |
| Total dose (e<sup>-</sup>/Å<sup>2</sup>) | 43 |  | 54.7 |  |
| Number of fractions | 30 |  | 40 |  |
| Defocus range (μm) | −0.5 to −2.9 |  | −0.5 to −2.9 |  |
| Number of micrographs | 12,797 |  | 23,966 |  |
| Acquisition software | Thermo Scientific EPU |  | Thermo Scientific EPU |  |
|  | **T = 3** | **T = 4** | **T = 3** | **T = 4** |
| EMDB ID | EMD-50832 | EMD-50833 | EMD-50834 | EMD-50835 |
| PDB ID | PDB-9FWE | PDB-9FWF | N/A | N/A |
| Number of particles contributing to map | 40,254 | 56,416 | 298,802 | 57,537 |
| Map resolution (FSC = 0.143) (Å) | 3.5 | 3.3 | 3.0 | 3.5 |
| Map resolution range around atom positions (Å) | 3.2 – 4.2 | 3.1 – 5.4 | 2.8 – 3.6 | 3.3 – 6.0 |
| Map sharpening B factor (Å<sup>2</sup>) | -187 | -179 | -203 | -223 |

